# Structural characterization of antibodies for synergistic HCMV neutralization

**DOI:** 10.64898/2026.08.22.746424

**Authors:** Artem Ashurov, Jory A. Goldsmith, Anna Ashurov, Zeyi Yu, Manuel Koch, Nica Classen, Linda Schlachter, Zahra Saidi, Danylo Shestopal, Katja Hoffmann, Hartmut Hengel, Vibor Laketa, Jason S. McLellan, Matthias Zehner, Florian Klein

**Author notes:** These authors contributed equally to this work (Artem Ashurov, Jory A. Goldsmith). Department of Cell and Virus Structure, Max Planck Institute of Biochemistry; Martinsried, 82152, Germany (Jory A. Goldsmith). These authors contributed equally to this work (Jason S. McLellan, Matthias Zehner, Florian Klein).

## Abstract

The limited mechanistic understanding of Human Cytomegalovirus (HCMV) infection and neutralization has hindered the effective application of antibodies for HCMV prevention and therapy. To address this, we conducted high-resolution cryo-EM analysis of 17 human monoclonal antibodies targeting HCMV glycoproteins gH and gL. This analysis revealed a landscape of vulnerable epitopes, including detailed binding modes, spatial orientations, and structural features associated with distinct neutralization phenotypes. By comprehensively characterizing the neutralizing properties of these antibodies and their combinations, we identified synergistic antibody cocktails that enhance the neutralization capacity of individual antibodies. Notably, while single antibodies only partially neutralized HCMV, their synergistic combinations achieved substantially more complete inhibition. Together, these findings delineate structural correlates of HCMV neutralization and establish functional principles for antibody-based combinatorial targeting of HCMV.

**One Sentence Summary:** This study offers an in-depth view of HCMV antibody neutralization, revealing critical target sites and synergy between antibody combinations.

## Introduction

Human cytomegalovirus (HCMV) is a ubiquitous pathogen capable of causing severe and fatal disease, particularly in immunocompromised individuals, fetuses, and newborns (*1*, *2*). Despite this threat, there is currently no licensed HCMV vaccine available and antiviral treatment options do not address all clinical needs (*3–5*). Antibody-based therapeutics have proven efficacious against multiple viral infections (*6–8*). However, the limited mechanistic insight into antibody responses against HCMV has hindered their effective application for prevention and therapy (*9*, *10*). To fully harness the potential of antibodies and advance efforts against HCMV, a deeper understanding of the structural and functional determinants underlying antibody-mediated neutralization is essential.

HCMV employs a sophisticated system involving multiple surface proteins to infect a broad range of host tissues and cells (*1*, *11*). In a previous study (*12*), we isolated a unique and potent set of human monoclonal antibodies (mAbs) targeting the glycoproteins H and L (gH/gL), which form the core of both the gH/gL/gO (trimer) and the gH/gL/UL128–131A (pentamer) complexes, critical components of HCMV infection machinery that facilitate host receptor binding and are involved in the membrane fusion during viral entry (*11*, *13*, *14*). Detailed epitope mapping revealed distinct antibody clusters, including previously undescribed binding sites on the gH/gL complex (*12*).

Here, we structurally and functionally characterized 17 representative mAbs from this panel, revealing novel epitope sites and identifying a consistent, epitope-specific synergy of antibody combinations that enhances the neutralization capacity and overcomes partial neutralization by individual mAbs. Together, these findings establish a structural and mechanistic framework to inform the development of antibody-based therapies and vaccines against HCMV.

## Results

### Competition-guided antibody selection enables simultaneous cryo-EM analysis

To define and characterize the novel binding sites of our previously identified 109 gH/gL mAbs (*12*), we assessed their epitope overlap with reference mAbs MSL-109, 3G16, 11B12, and 13H11 (*15*, *16*), which recognize two known gH epitopes, site A and site B (*17*) (**Fig. 1A**, left). Based on competition patterns, we grouped our mAbs into those targeting site A or B and additional, previously undefined epitope groups (I–VI) (*12*) (**Fig. 1A**, right). Next, we selected unique mAbs representative of each gH/gL competition group and designed two combinations of non-overlapping mAbs for subsequent analyses, based on ELISA competition experiments performed in a mAb-vs-mAb setting (**Fig. 1B**). Selection 1 consisted of nine mAbs, and Selection 2 included eight mAbs (**Fig. 1B, fig. S1A**). Both selections encompassed, wherever possible, the most potent neutralizing mAbs for each competition group with IC_50_ values as low as 0.02 µg/ml and up to 100% neutralization capacity in fibroblast cells (**fig. S1B, fig. S1C**). To further evaluate the selected mAbs, we tested their neutralization performance against a panel of HCMV-susceptible host cells, including fibroblast-like cells and epithelial-like cells (**Fig. 1C**). While some mAbs (e.g. site VI_3_ mAbs) were mostly non-neutralizing, others consistently demonstrated low IC_50_ values (<0.1 µg/ml) across most tested cell lines and achieved up to 100% neutralization capacity.

**Fig. 1.**
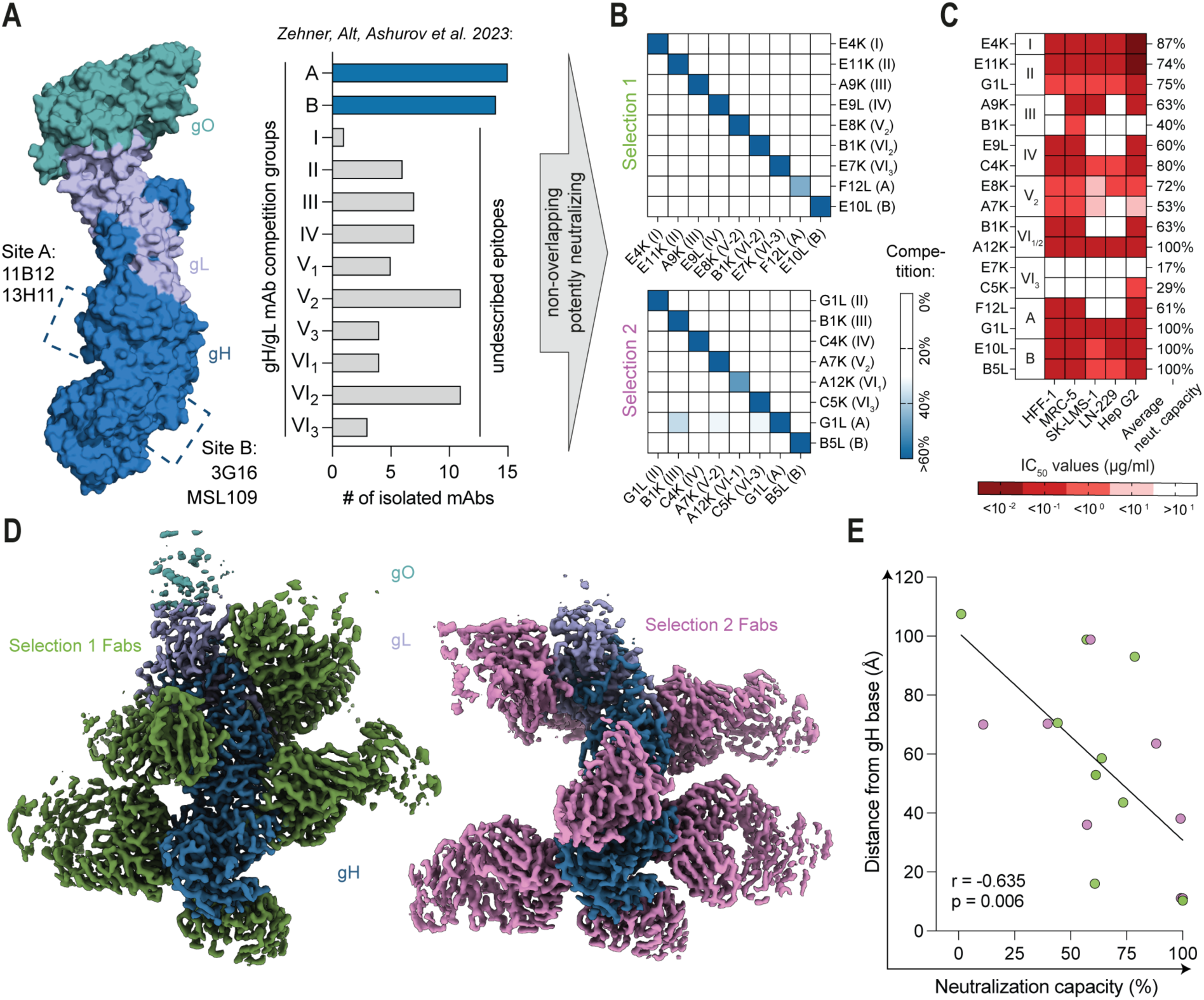
Characterization of antibodies selected for simultaneous cryo-EM. (**A**) Overview of HCMV gH/gL-targeting mAbs. Left: MSL-109, 3G16, 11B12, and 13H11 with corresponding binding sites. Right: Number of recently isolated mAbs against each competition group, with additional groups with undescribed binding sites displayed in gray. (**B**) Competition among selected mAbs within each selection in a mAb-vs-mAb setting, determined by ELISA. Competition activity is highlighted in blue. (**C**) Neutralizing performance of each mAb in blocking HCMV_TB40/E_ infection across various cell lines, represented as IC_50_ values (µg/ml) in sequential red colors. The average neutralization capacity for each mAb across all tested cell lines is shown on the right (100% indicating complete neutralization at maximal tested antibody concentration). (**D**) Composite cryo-EM maps of Fabs corresponding to each selection of mAbs in complex with HCMV trimer. Left: Selection 1. Right: Selection 2. (**E**) Correlation between the neutralization capacity of each mAb (x-axis, data from **fig. S1C**) and the distance of its binding site from the gH base (y-axis). Distances were measured from a plane approximating the bottom of the gH base to the center of the Fab epitope footprint. Correlation was assessed using the Pearson correlation coefficient (r), with a straight line representing a simple linear regression model.

To identify the molecular epitopes recognized by the selected mAbs, we determined 3.2Å-resolution cryo-EM structures of the HCMV trimer bound simultaneously to 9 and 8 Fab fragments from Selections 1 and 2, respectively (**Fig. 1D**, **fig. S1D-G**). 14 of the 17 Fabs in the two reconstructions were resolved to high-resolution, enabling model building and identification; the remaining three Fabs showed only weak density in the maps but were identifiable based on their competition profiles. Interestingly, we identified an inverse correlation between mAb neutralization capacity and the distance of the epitope from the gH base, with all mAbs that achieved complete HCMV neutralization binding membrane-proximal regions (**Fig. 1E**).

Together, these findings establish a structural framework linking epitope location on gH/gL to neutralization capacity of targeting mAbs.

### Structural mapping defines distinct vulnerable epitopes on the gH/gL complex

We found that 8 of the 17 Fabs bound to the gH base, which arise from competition groups A (F12L, G1L), B (E10L, B5L), V_2_ (E8K, A7K), and VI_1/2_ (A12K, B1K; these Fabs compete despite being grouped in VI_1_ and VI_2_, respectively) (**Fig. 2**). Antibodies in group A compete with 13H11, and antibodies in group B, and to a small extent, antibodies at site VI_1/2_, compete with MSL-109 (**Fig, 2,** right), both of which have been structurally characterized at high resolution (*13*).

**Fig. 2.**
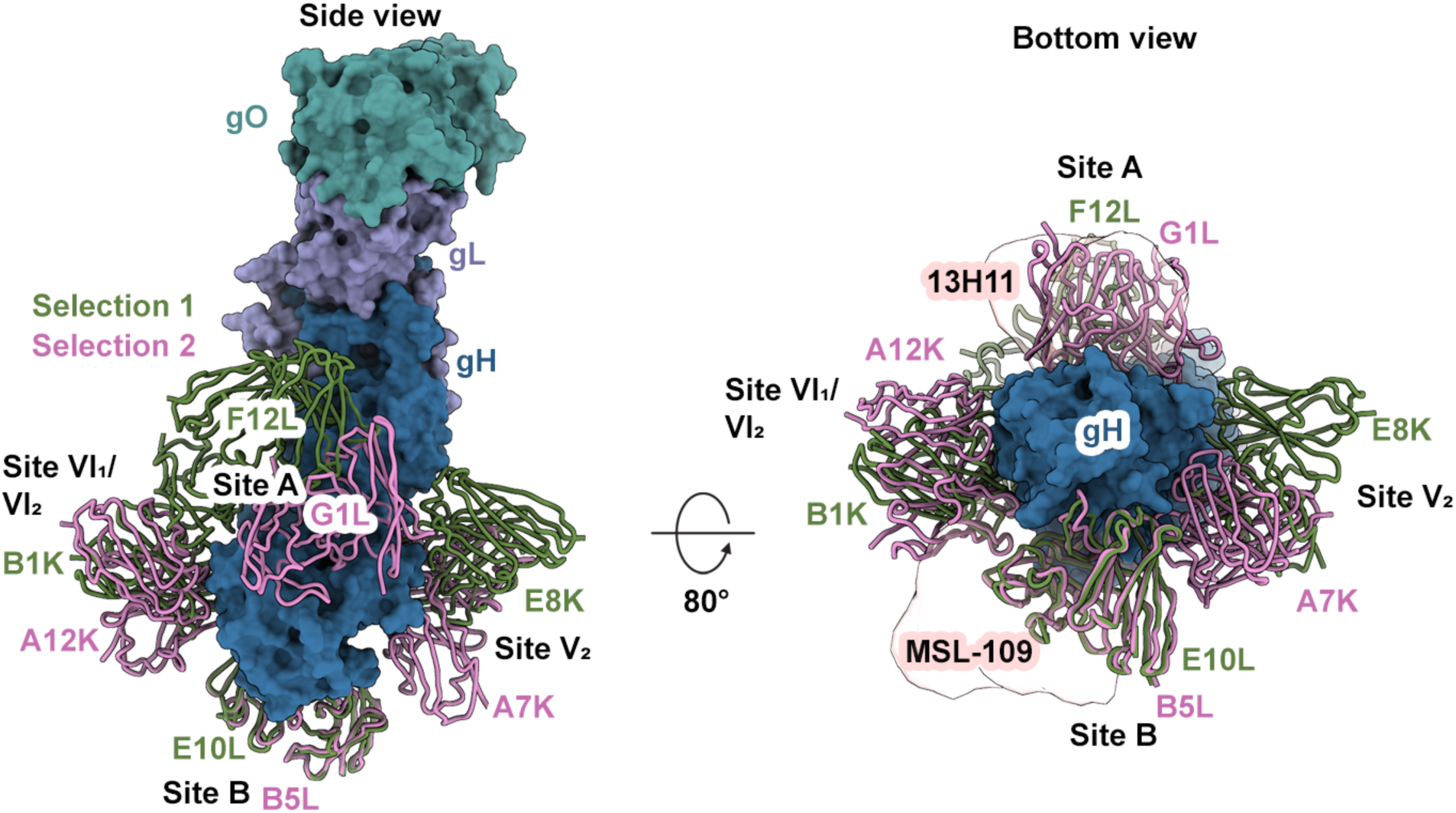
Structures of gH base-binding antibodies. Left: Side view of gH base-binding Fabs F12L, E8K, E10L, B1K (green, Selection 1) and G1L, A7K, B5L, and A12K (pink, Selection 2). Fabs are shown as backbone traces and HCMV trimer is shown as a molecular surface with gH in dark blue, gL shown in purple, and gO shown in teal. Right: bottom view of the same Fabs bound to the gH base, with transparent peach molecular surfaces showing published antibodies 13H11 and MSL-109.

Inspection of group A antibodies reveals that F12L, G1L, and reference mAb 13H11 each have different epitopes that only overlap partially with one another (**Fig. 2**, right). G1L binds closer to the membrane on the same face of gH and inserts into a pocket formed by DII, DIII, and DIV, primarily interacting with DIII via its CDRH3. By contrast, F12L targets a more membrane-distal region of DII than reference mAb 13H11 in gH α2, which is part of the region that connects DII to DI and wraps around gL as part of the gH-gL interface (**fig. S2A**). Group B antibodies E10L and B5L bind the same epitope in the gH heel (**Fig. 2**). Compared to the reference mAb MSL-109, which contacts gH DIV but primarily interacts with DIII, E10L and B5L mainly recognize DIV and are more membrane-proximal (**fig. S2B**). Antibodies in groups A and B bind to opposing faces of gH/gL, while antibodies in groups V_2_ and VI_1/2_ engage the remaining empty faces that are approximately perpendicular to the groups A and B epitopes (**Fig. 2**, right). Specifically, group V_2_ epitopes are approximately 90 degrees clockwise from group A epitopes (looking towards the membrane) and group VI_1/2_ are approximately 90 degrees counterclockwise from group A epitopes (**Fig. 2**, right). Group V_2_ antibodies E8K and A7K bind at the same site, however the angle of approach of A7K is rotated 90 degrees relative to E8K such that it targets a different set of residues. Whereas E8K primarily binds helices in DII, A7K contacts part of the gH coil region from residues 147-161 and DIII helices (**fig. S2C**). Lastly, the group VI_1/2_ antibodies B1K and A12K bind the same site in gH DIII and DIV. However, B1K has an angle of approach that is rotated 90 degrees relative to A12K, allowing it to contact a more membrane-distal part of gH DIII (**fig. S2D**). Notably, A12K fully neutralizes HCMV, whereas B1K neutralizes it to only 63% (**Fig. 1C**), despite binding to almost all of the A12K epitope residues. This suggests that the angle of approach, which positions the A12K Fab more proximally to the membrane, may be crucial for its higher neutralization capacity at this site.

The remaining 9 antibodies bound to more membrane-distal epitopes around the gH-gL interface (**Fig. 3**). 3 of the 17 antibodies were poorly resolved in the 2 cryo-EM maps and therefore could not be identified based on sequence. However, a low-resolution Fab smear is present in both maps around the gH N-terminus, suggesting they are competing Fabs E11K and G1L (group II) (**Fig. 3B**). These Fabs likely recognize a flexible portion of the gH N-terminus as they are too far from any rigid part of the trimer. E4K (group I) binds at the gH/gL interface at a similar height and angle to the group II antibodies at the gH N-terminus, and directly clashes with cellular host receptor TGFBR3 (**fig. S3A**). A9K and B1K (group III) target a more membrane-proximal site on the gH/gL interface consisting of the gL C-terminal region and gH DII. B1K binds more membrane-proximally and also points its Fc more toward the membrane (**Fig. 3A**). A9K and B1K binding to pentamer could inhibit THBD-induced pentamer dimerization (*12*, *18*) by clashing with the adjacent pentamer (**fig. S3B**).

**Fig. 3.**
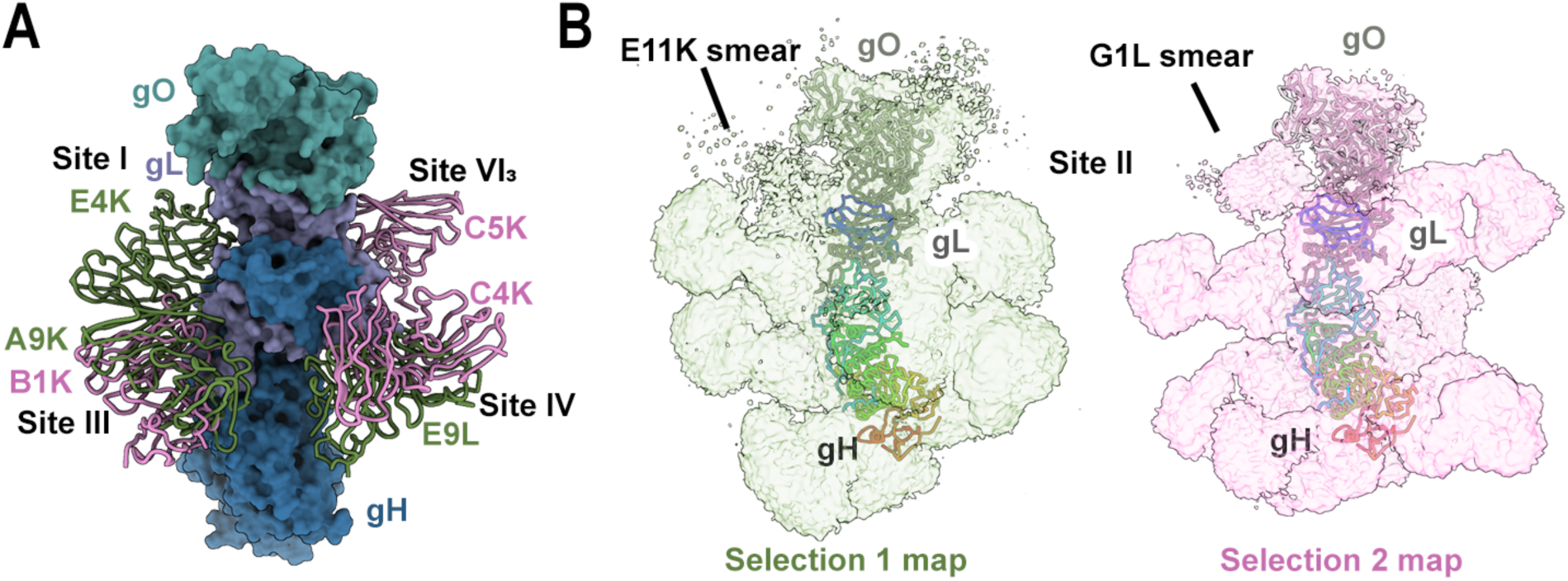
Structures of membrane-distal gH/gL antibodies. (**A**) Side view of membrane-distal gH/gL Fabs E9L, A9K, E4K (green, Selection 1), and C5K, C4K, and B1K (pink, Selection 2). Fabs are shown as backbone traces and HCMV trimer is shown as a molecular surface with gH in dark blue, gL shown in purple, and gO shown in teal. (**B**) Low threshold maps for Selections 1 and 2 showing smeared Fab density at Site II for E11K (left, Selection 1) and G1L (right, Selection 2).

Interestingly, C4K (group IV) bound an almost identical epitope to our recently published neutralizing antibody CS4tt1p1_E3K (referred to as E3K) (*12*), and contacts a gL segment containing α1 as well as gH DI. However, the competing group IV antibody, E9L, binds the same site but only contacts the gL α1-containing segment and forms almost no contacts with gH (**fig. S3C**). E3K and C4K exhibit higher neutralization capacity than E9L, suggesting that engagement of gH may contribute to enhanced neutralization at this site. Notably, C5K (group VI_3_) binds to a gL segment just downstream of α1 and also does not contact gH (**fig. S3D**). C5K shows very low neutralization capacity (**Fig. 1C**), suggesting a mechanism similar to that of group IV antibodies.

Lastly, the epitope of the third low-resolution Fab (E7K) was only discovered upon 3D classification of the reconstruction for Selection 1, which revealed one sub-class containing gO and no E7K, and one sub-class where gO was unresolved and was replaced by a Fab binding to the top of gL (**fig. S3E**). Therefore, E7K appears to displace gO or render it more flexible to accommodate the clash. In addition, the position of E7K results in a steric clash in 3D with C5K (**fig. S3F**).

In summary, our cryo-EM structures define a spatially organized epitope landscape, revealing novel and distinct sites of vulnerability across HCMV gH and gL.

### Antibodies act synergistically to enhance HCMV neutralization

We next evaluated the neutralizing activity of different mAb combinations by pairing antibodies in a 1:1 ratio and analyzing the performance of the resulting cocktails. Interestingly, while combining two potent mAbs did not result in significant additive effects on IC_50_ values (**fig. S4A**), certain pairings of weaker mAbs produced potent neutralizing cocktails (e.g., antibodies targeting site III paired with C5K mAb of site VI_3_) (**fig. S4A**). Individually, these mAbs displayed only partial neutralizing activity, plateauing at high levels of residual infection even at increasing antibody concentrations (hereafter referred to as ‘high-plateau’ mAbs) (**Fig. 4A**, **fig. S1B**). However, when combined, these mAbs achieved near-complete neutralization, reaching up to 90–100% neutralization capacity (**Fig. 4A**).

**Fig. 4.**
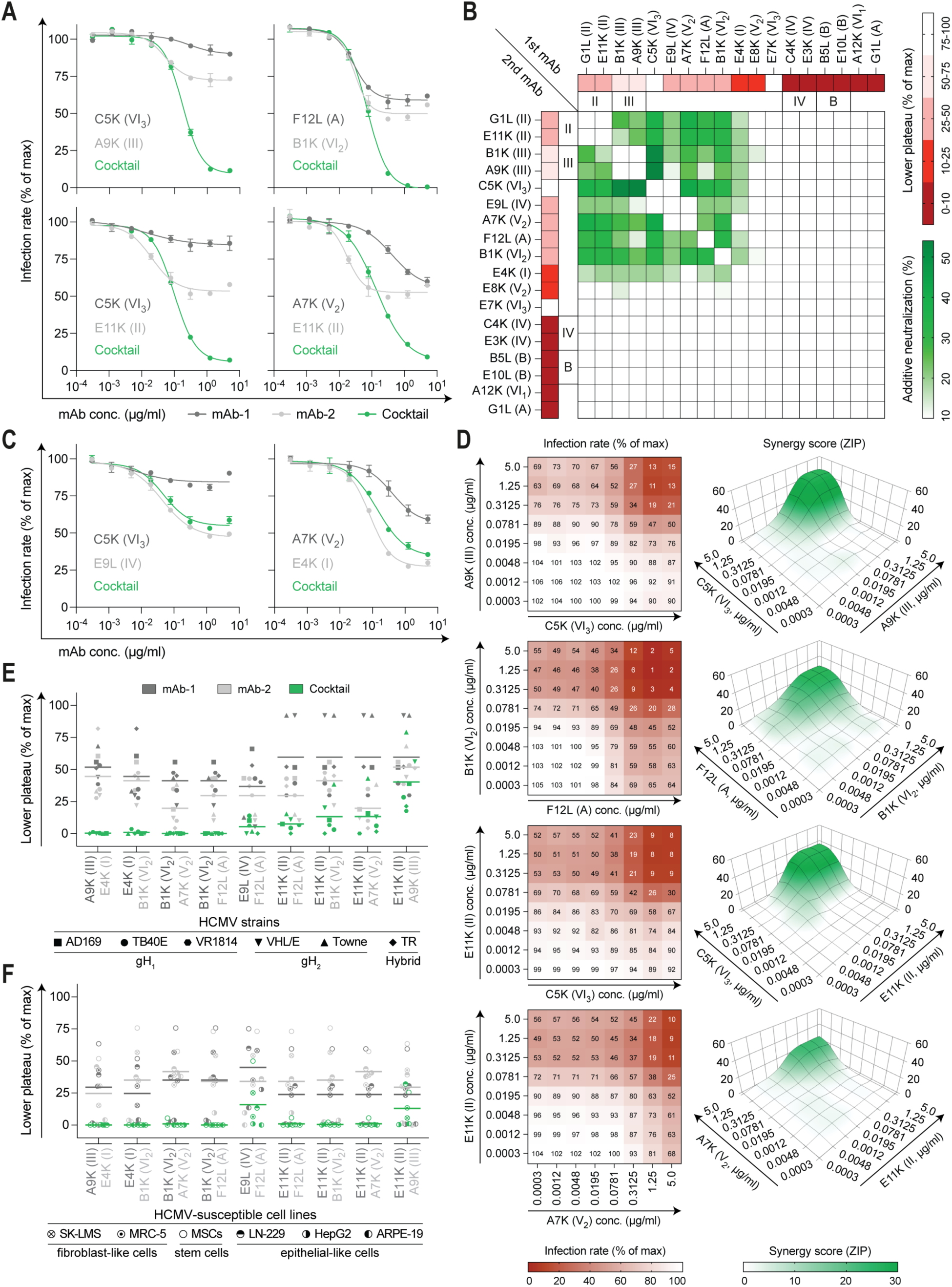
Synergistic action of antibodies in neutralizing HCMV. (**A**) Neutralizing activity of individual mAbs and their cocktails against HCMV_TB40/E_ infection in HFF-1 fibroblasts. The mAb with the lowest neutralization capacity is shown in dark gray, the second mAb in light gray. Cocktails of these mAbs are displayed in green on the same graph. Infection rates are expressed as percentages relative to the no-antibody (maximal infection) control. Mean values and SDs from two separate experiments are shown. (**B**) Synergistic effects (displayed in green) for all possible combinations of the mAb panel (primary data from **fig. S4B**). Lower plateau values (i.e. residual infection levels) of single mAbs are shown in shades of red. Additive neutralization is measured as the difference between the lower plateau level of a cocktail and the lowest plateau of individual mAbs. The graph is mirrored along the main diagonal. mAbs are clustered by additive neutralization levels and epitope representation. ‘High-plateau’ mAbs and their combinations were tested in at least two separate experiments; representative data are shown. (**C**) Neutralizing activity of individual antibodies and their non-synergistic cocktails against HCMV_TB40/E_ infection in HFF-1 fibroblasts. The mAb with the lowest neutralization capacity is shown in dark gray, the second mAb in light gray. Cocktails of these mAbs are displayed in green on the same graph. Mean values and SDs from two separate experiments are shown. (**D**) Neutralizing activity of cocktails with different stoichiometries assessed by testing varying concentrations of cocktail members against HCMV_TB40/E_ infection in HFF-1 fibroblasts. Left: Percentage of residual infection achieved by each concentration. Mean values from two separate experiments are displayed. Right: 3D ZIP (Zero Interaction Potency) synergy landscapes for indicated antibody combinations across dose matrices. Surfaces show ZIP synergy scores (z-axis) over the tested concentration ranges of each antibody in the combination (x- and y-axes, respectively). Higher ZIP synergy scores are shown in darker green. Positive scores suggest an additive interaction, whereas a score greater than 10 indicates a synergistic interaction. Displayed surfaces were created and analyzed using the SynergyFinder tool (https://synergyfinder.org) (*19*). (**E**) Lower plateau levels of individual mAbs and their cocktails against different HCMV strains against HFF-1 fibroblast infection. Different gH genotypes within HCMV strains are noted below. (**F**) Lower plateau levels of individual mAbs and their cocktails against HCMV_TB40/E_ infection in different host cell lines. (**E-F**) Each dot represents the mean from two separate experiments. Lines indicate arithmetic means for each antibody/cocktail.

Importantly, this synergistic effect was observed for multiple gH/gL antibody combinations across different epitopes, with most cocktails of ‘high-plateau’ mAbs exhibiting enhanced neutralization through a reduction of the residual infection plateau (**Fig. 4A-B**, **fig. S4B**). Antibodies recognizing the same target site showed no synergy when combined (e.g., group II antibody combination) (**Fig. 4B**). Notably, some combinations of ‘high-plateau’ mAbs targeting different epitopes were non-synergistic – e. g., C5K (VI_3_) with E9L (IV) (**Fig. 4C**, **fig. S4C**) – even though each of these mAbs demonstrated synergy when combined with other ‘high-plateau’ mAbs (**Fig. 4B**). This observation aligns with structural data, which suggest that C5K and E9L likely share a similar neutralization mechanism (**fig. S3C-D**), indicating that synergistic neutralization requires antibodies with distinct modes of action.

We then investigated the optimal stoichiometry of the cocktails by testing different ratios of their components, confirming that synergy was best maintained at a 1:1 ratio (**Fig. 4D**, left). We next quantified these effects using the ZIP (Zero Interaction Potency) model, in which scores greater than 10 are considered indicative of synergy. This analysis demonstrated robust synergy across all four tested mAb combinations (**Fig. 4D**, right). The strongest effect was observed for the C5K/A9K cocktail, which reached a mean ZIP score of 13.17 across the dose matrix and a maximal ZIP score of 48.91 (**Fig. 4D**, right).

To assess whether this effect is conserved against different HCMV strains, we selected nine cocktails across different epitopes and tested their performance against a panel of six strains representing distinct HCMV gH genotypes (**Fig. 4E**). We found that synergy was consistently observed across all tested strains, with single mAbs showing weaker neutralization capacity compared to their cocktails (**Fig. 4E**). To validate the synergy findings, we tested six additional HCMV-susceptible host cell lines (**Fig. 4F**). Similarly, cocktails outperformed single mAbs in terms of lowering the plateau. Notably, most of ‘high-plateau’ mAbs were able to neutralize HCMV with up to 100% capacity in some epithelial-like cells (HepG2 and ARPE-19) (**Fig. 4F**, **fig. S4D**).

Collectively, these data demonstrate epitope-dependent antibody synergy that reduces residual infection plateaus, is conserved across HCMV strains and host cell types, and enables enhanced neutralization through combinatorial targeting. Together with our structural analyses defining novel vulnerable epitopes on the gH/gL complex, these findings establish functional principles for rational antibody-based intervention strategies.

## Discussion

To date, the only approved antibody-based product used clinically in the HCMV setting is Cytogam, a HCMV hyperimmune globulin preparation derived from human plasma with anti-HCMV antibody content (*20*). Although Cytogam established proof of principle for antibody-based prophylaxis (*20–22*), it also highlights key drawbacks of existing approaches, as it is a heterogeneous polyclonal product with suboptimal potency (*23*, *24*). These limitations underscore the need for next-generation antibody-based strategies, including defined monoclonal antibody (mAb) approaches. However, major obstacles to advancing such approaches include the limited availability of highly potent neutralizing mAbs and incomplete understanding of the underlying neutralization determinants (*9*, *10*, *25*).

Using cryo-EM, we performed detailed structural characterization of 17 novel HCMV-targeting human mAbs, uncovering previously undescribed molecular epitopes on the gH/gL complex. These epitopes are organized into two structural tiers — the gH base and the gH/gL interface — forming distinct epitope belts that can associate with distinct neutralization phenotypes. This structural framework provides a foundation for mechanism- and epitope-guided mAb selection and the rational design of optimized antibody combinations for further preclinical and clinical evaluation in therapeutic and prophylactic settings.

Notably, we found that mAbs capable of complete HCMV neutralization bound near the gH base, while those targeting membrane-distal regions consistently exhibited lower neutralizing capacity. These insights may guide vaccine strategies designed to focus immune responses toward vulnerable sites on gH/gL, in particular toward eliciting gH base-specific antibodies with high neutralization potential.

Evidence from clinical trials suggests that combining antibodies, such as in the RG7667 (*26*) or CSJ148 (*27*) cocktails, can improve clinical outcomes. Although neither trial met its primary endpoint, both demonstrated antiviral activity, including reduced viral load or a lower incidence of HCMV disease (*26*, *27*). Our findings reinforce the potential of antibody combinations in addressing the limitations of single-antibody therapies. Our data are consistent with a model in which binding of a single ‘high-plateau’ mAb fails to fully disable gH/gL function, leaving the complex sufficiently active to mediate infection through alternative pathways, permitting residual viral entry despite high-affinity engagement. This incomplete inhibition may explain the residual infection observed with individual mAbs. Importantly, we show that synergistic cocktails can prevent this residual infection and achieve more complete inhibition of HCMV through combinatorial targeting of non-overlapping epitopes, likely by simultaneously targeting multiple functional sites on the gH/gL complex. Together, these findings highlight a set of functional principles for antibody-based intervention strategies against HCMV.

### Limitations of the study

This study has limitations that warrant consideration. First, the selected panel of mAbs may not encompass the full diversity of gH/gL-targeting antibodies, as our selection was based on observed competition profiles and prioritization of mAbs with potent neutralizing activity within each epitope group. Additionally, we focused on high-affinity mAbs with comparable ELISA EC_50_ values, potentially excluding functionally relevant but lower-affinity mAbs that may contribute to polyclonal responses in vivo.

Second, our investigation of synergistic effects was limited to gH/gL-specific mAbs due to the absence of reported ‘high-plateau’ mAbs targeting other HCMV glycoproteins. However, synergy may be a broader phenomenon applicable to other antigenic targets.

Finally, while the synergistic activity of mAb combinations was demonstrated in vitro, in vivo validation is needed to determine whether such combinations can achieve superior HCMV neutralization compared to individual mAbs and to assess their therapeutic potential under physiological conditions.

## Materials and Methods

### Study design

This study combined biochemical, structural, and functional approaches to define the basis of antibody-mediated neutralization of human cytomegalovirus (HCMV). Binding and competition ELISAs were first used to map epitope relationships and identify non-overlapping human monoclonal antibodies targeting distinct regions of the HCMV gH/gL complex. These data enabled the rational design of two antibody groups for simultaneous high-resolution cryo-EM analysis, allowing 17 monoclonal antibodies to be resolved in only two gH/gL/gO–antibody cryo-EM structures. Antibodies were then functionally characterized alone and in combination in cell culture-based neutralization assays to identify synergistic interactions and measure their effects on neutralization potency and residual infection levels. Together, these approaches enabled structural definition of neutralizing epitopes and functional identification of antibody combinations with enhanced antiviral activity.

### Bacterial Strains

Escherichia coli DH5α strain (Thermo Fisher Scientific) was used for plasmid amplification of previously cloned expression vectors of HCMV-targeting monoclonal antibodies (*12*).

### Cells and viruses

Primary HFF-1 fibroblast cells (ATCC) were maintained in MEM supplemented with GlutaMAX™ (Gibco), 10% FBS (Sigma-Aldrich), 0.1 mg/mL gentamicin (Sigma-Aldrich) and 1 µg/L Human FGF-basic Recombinant Protein (Gibco). MRC-5, SK-LMS-1, LN-229 and HepG2 cell lines (ATCC) were maintained in DMEM supplemented with L-Glutamine (Gibco), sodium pyruvate (Sigma-Aldrich), 10% FBS and 0.1 mg/mL gentamicin. ARPE-19 cells (ATCC) were maintained in DMEM / F12 (1:1) supplemented with L-Glutamine, 15 mM HEPES (Gibco), 10% FBS and 0.1 mg/mL gentamicin. Bone marrow-derived mesenchymal stem cells (MSCs; ATCC-PCS-500-012) were maintained in Mesenchymal Stem Cell Basal Medium (ATCC) supplemented with Mesenchymal Stem Cell Growth Kit for Bone Marrow-derived MSCs (ATCC) and 0.1 mg/mL gentamicin.

Virus stocks were generated by propagating replication-competent HCMV strains in HFF-1 cells for two consecutive 7-day passages, with supernatant transfer to fresh cells between passages. Cell-free supernatants were subsequently harvested, aliquoted, and stored at –80°C for use in neutralization assays. Each virus stock was titrated via fluorescence detection of HCMV immediate early antigen 1/2 (IE1/2; see staining protocol below) to determine infectious concentration. The following HCMV strains were used: HCMV_TB40/E_ (*28*), HCMV_AD169_ (*29*), HCMV_Towne_ (*30*), HCMV_VHL/E_ (*31*), HCMV_VR1814_ (*32*), and HCMV_TR_ (*33*).

### HCMV glycoprotein production and purification

For binding ELISA, the Sleeping Beauty transposon system was applied to generate the gH/gL/gO (trimer) protein, as previously described (*12*). Three separate constructs included: gH (Uniprot P12824; HCMV strain AD169; AA 27–720; BM40 signal peptide; C-terminal Twin-Strep-tag), gL (Uniprot F5HCH8; HCMV strain Merlin; AA 32–278; BM40 signal peptide), and gO (Uniprot P16750; HCMV strain AD169; AA 40–466; BM40 signal peptide). Recombinant protein production was performed by co-transfecting the expression constructs and the transposase plasmid at a 10:1 ratio into HEK293 EBNA cells using FuGENE® HD transfection reagent (Promega GmbH, Madison, USA). Post-transfection, cells underwent stringent puromycin selection (2 µg/mL; Sigma), were expanded in triple flasks, and protein expression was induced with doxycycline (0.5 µg/mL; Sigma). Supernatants from confluent cultures were filtered and purified using Strep-Tactin® XT resin (IBA Lifescience, Göttingen, Germany). Purified recombinant HCMV proteins were eluted using biotin-containing buffer (IBA Lifescience), dialyzed against PBS, and aliquots stored at −80°C.

For antibody-competition ELISA, the pentamer complex (gH/gL/UL128–131A) was purchased from ‘The Native Antigen Company’, aliquoted and stored by −80°C.

For structural analyses, plasmids encoding HCMV gO, HCMV gL, and HCMV gH-His ectodomain were transiently co-transfected into Freestyle 293F cells using polyethylenimine (Polysciences), and cells were incubated for 6 days at 37°C after which the supernatant was harvested. HCMV trimer was purified from the supernatant with NiNTA agarose and ran over a Superdex 200 10/300 GL (Cytiva).

### In vitro production and purification of monoclonal antibodies

The sequences of monoclonal antibodies used in this study were previously reported (*12*), and the antibody production followed established protocol (*34*). In brief, HEK293-6E cells (National Research Council Canada) were transfected using branched polyethylenimine (PEI, Sigma-Aldrich). For transfection, 50 mL cultures of HEK293-6E cells at a density of 0.8×10^6^ cells/mL were co-transfected with 25.0 µg of heavy chain plasmid and 25.0 µg of light chain plasmid. Cells were cultured in FreeStyle 293 Expression Medium (Thermo Fisher Scientific) supplemented with 0.2% Penicillin/Streptomycin (Thermo Fisher Scientific) at 37°C in 6% CO_2_. Seven days post-transfection, supernatants were collected and filtered through 0.2 µm Nalgene Rapid-Flow filters (Thermo Fisher Scientific). Filtered supernatants were incubated with Protein G Sepharose 4 Fast Flow beads (Sigma-Aldrich) overnight at 4°C. The beads with bound antibodies were subsequently washed in 10 ml chromatography columns (BioRad), and antibodies were eluted using 0.1 M glycine (pH 3), and immediately buffered with 1 M Tris (pH 8). Finally, a buffer exchange to PBS was performed using 30 kDa Amicon Ultra-4 spin columns (Millipore), and the purified antibodies were stored at 4°C.

### ELISA for antibody binding activity

For binding ELISA, 96-well High Bind microplates (Corning) were coated overnight at 4 °C with 3 µg/ml of trimer in PBS. Plates were then washed three times with PBST (PBS containing 0.05% Tween-20; Carl Roth) and blocked with PBST supplemented with 5% nonfat dried milk powder (PanReac AppliChem) for 60 minutes at room temperature (RT). gH/gL-specific antibodies were added at a starting concentration of 30 µg/ml in serial 5-fold dilutions and incubated with the coated protein for 90 minutes at RT. After another three washes, plates were incubated with horseradish peroxidase (HRP)-conjugated goat anti-human IgG secondary antibody (SouthernBiotech, 2040-05; 1:2000 in PBST with 5% milk powder) for 45 minutes at RT. Signal development was performed using ABTS substrate (2,2’-azino-bis(3-ethylbenzothiazoline-6-sulfonic acid); Thermo Fisher Scientific), and absorbance was measured at λ_415-695_ nm using a Tecan plate reader after a fixed incubation period. Binding curves and EC_50_ values were generated ana analyzed using GraphPad Prism 9.

### Antibody-competition ELISA

Prior to the competition ELISA, 0.5 mg of each antibody was biotinylated with a 50x molar excess of biotin using the EZ-Link Sulfo-NHS-Biotin (Thermo Fisher) as per manufacturer’s protocol, and a buffer exchange to PBS was performed using 30 kDa Amicon Ultra-4 spin columns (Millipore).

For antibody competition ELISA, 96-well High Bind microplates (Corning) were coated overnight at 4 °C with 3 µg/ml of pentamer in PBS. Plates were then washed three times with PBST (PBS containing 0.05% Tween-20; Carl Roth) and blocked with PBST supplemented with 5% nonfat dried milk powder (PanReac AppliChem) for 60 minutes at RT. Unbiotinylated gH/gL-specific antibodies were added at a concentration of 30 µg/ml and incubated with the coated protein for 90 minutes at RT. After another three washes, biotinylated antibodies were added at their EC_50_ concentrations and incubated for 45 minutes at RT. Then, plates were incubated with horseradish peroxidase-conjugated Streptavidin (Jackson ImmunoResearch, 1:5000 in PBST / 5% Nonfat dried milk powder) for 30 min at RT. Signal development was performed using ABTS substrate (2,2’-azino-bis(3-ethylbenzothiazoline-6-sulfonic acid); Thermo Fisher Scientific), and absorbance was measured at λ_415-695_ nm using a Tecan plate reader after a fixed incubation period. Optical density values were normalized to biotinylated-antibody-only control to calculate percent competition. The corresponding unbiotinylated antibody was included in each experiment to verify maximum competition. Competition heatmaps were generated and analyzed using GraphPad Prism 9.

### Virus neutralization tests

For evaluation of neutralizing activity of mAbs and their combinations against HCMV infection in HFF-1 fibroblasts, 96-well Clear Flat Bottom TC-treated microplates (Corning) were seeded with 12000 cells/well in MEM supplemented with GlutaMAX™ (Gibco), 10% FBS (Sigma-Aldrich) and 0.1 mg/mL gentamicin (Sigma-Aldrich) (referred to as HFF-NT medium) and incubated overnight at 37°C. The following day, antibody samples were serially diluted in HFF-NT medium, starting at a concentration of 10 µg/mL. Diluted antibodies were mixed 1:1 (final starting concentration of antibodies = 5 µg/mL) with HCMV_TB40/E_ or other virus stock at the multiplicity of infection (MOI) of 1 infectious unit per cell or below (MOI of approximately 0.5–1.0), and preincubated for 2 hours at 37°C. Subsequently, the virus-antibody mixtures were added to HFF-1 cells and incubated for 18-20 hours at 37°C. Then, the cells were fixed with 80% acetone (Carl Roth) for 5 minutes at RT and washed three times with PBST (PBS containing 0.05% Tween-20; Carl Roth). Next, plates were stained with mouse mAb targeting HCMV immediate early antigen 1/2 (1:1000 in PBS; clone CH160; Abcam) overnight at 37°C. After three washes with PBST, plates were stained with Cy3-AffiniPure F(ab’)2 Fragment Goat Anti-Mouse IgG (H+L) (1:300 in PBS; Jackson ImmunoResearch) and 1 µg/mL DAPI (Miltenyi Biotec) for 1 hour at 37°C, and then washed three times again.

The fluorescence-readout of stained plates was done using the ImageXpress Pico system, quantifying the number of infected cells relative to the total number of cells. Infection rates were expressed as percentages relative to the no-antibody (maximal infection) control. Dose-response curves generated from these data were subjected to non-linear regression analysis (using GraphPad Prism 9) to assess the neutralizing capacity (%) and neutralizing potency (IC_50_ values, µg/mL) of each antibody. Neutralizing capacity was defined as the difference between 100% infection (no-antibody control) and the lower plateau (minimum) of the dose-response curve. For example, if an antibody produced a lower plateau of 20% residual infection at saturating concentrations, its neutralizing capacity was considered 80%. If no clear lower plateau was reached at 5 µg/mL, the residual infection at that concentration was used to calculate maximal neutralizing capacity. IC_50_ values were calculated only for antibodies with a neutralizing capacity of at least 50%, and correspond to the antibody concentration at which 50% of the maximal neutralization effect was achieved (relative IC_50_) (*35*).

For testing neutralizing activity of mAbs in other HCMV-susceptible host cell lines, the protocol was modified as follows: For fibroblast-like cells (MRC-5 and SK-LMS-1), and epithelial-like cells (LN-229 and HepG2), DMEM supplemented with L-Glutamine (Gibco), sodium pyruvate (Sigma-Aldrich), 10% FBS and 0.1 mg/mL gentamicin was used. For ARPE-19 epithelial cells, DMEM / F12 (1:1) supplemented with L-Glutamine, 15 mM HEPES (Gibco), 10% FBS and 0.1 mg/mL gentamicin was used. For bone marrow-derived mesenchymal stem cells (MSCs), MEM supplemented with GlutaMAX™ (Gibco), 10% FBS and 0.1 mg/mL gentamicin was used. As epithelial-like cells (LN-229, HepG2, ARPE-19) are less susceptible to HCMV infection than fibroblast cells, 4- to 32-fold higher concentrations of virus were needed in those cell types as compared to HFF-1 fibroblasts to achieve absolute infection rates of 20-40% (MOI ≤ 0.5).

### Generation of Fab fragments from IgG for structural analyses

1 mg/ml of each IgG was incubated with 1:2000 wt/wt Lys-C endoproteinase (Thermo Fisher) for 24h at 37 °C. Lys-C reactions were quenched by the addition of 1X Roche cOmplete protease inhibitor cocktail (Sigma-Aldrich). Fabs were then passed over CaptureSelect CH1-XL Affinity Matrix (Thermo Fisher) and the Fc was allowed to flow through. Fabs were eluted with 0.1M glycine pH 3 and neutralized with 0.1M Tris pH 8. Fabs were then concentrated and flash-frozen.

### Cryo-electron microscopy

0.25 mg/ml HCMV trimer was mixed with a 2-fold molar excess of nine Selection 1 Fabs or a 2-fold molar excess of eight Selection 2 Fabs, and incubated for 30 min at room temperature. 3 μl of sample was then added to UltrAuFoil holey gold grids (Electron Microscopy Sciences) and blotted in a Vitrobot Mark IV (Thermo Fisher) using a blot time of 4 seconds, blot force of 0, 100% humidity at 4 °C. The Selection 1 and Selection 2 grids were imaged in a Glacios (Thermo Fisher), and 2979 and 2529 movies were recorded for each, respectively, at 30° tilt, 150,000x magnification (0.94 Å/px) and with defocus varying from −0.8 μm to −2.2 μm. A full description of the data collection statistics can be found in **Table S1**. Data collection was automated using SerialEM (*36*). Patch motion correction, patch CTF Estimation, blob picking, 2D classification, ab initio reconstruction, heterogeneous refinement, non-uniform refinement, and 3D classification were performed in cryoSPARC v3.3.2 (*37*). SAbPred ABodyBuilder (*38*) was used to generate Fab models for each Fab, which were then placed into the maps when they could be identified. Model-building was then performed in Coot (*39*) and ISOLDE (*40*), and refinement was performed in Phenix (*41*). Structural biology applications used in this project were compiled and configured by SBGrid (*42*).

### Quantification and statistical analysis

Quantitative and statistical analyses were performed using GraphPad Prism 9 and Microsoft Excel for Mac (v16.44). Microscopic analyses were done by ImageXpress Pico and Nikon Ti2 microscope. For structural analyses cryoSPARC, Phenix, Coot, ISOLDE, UCSF ChimeraX (*43*) and PyMOL were used.

## Acknowledgments

We gratefully thank Christian Sinzger as well as all members of the Koch, Laketa, Hengel, Klein, Zehner, and McLellan Laboratories for helpful discussion and support. Moreover, we thank Anna Schmitt, Tina Bresser, and Lisa Kottege for lab management and assistance.

## Funding

German Center for Infection Research TTU 07.835 (FK)

German Center for Infection Research MD Program TI 07.003 (Artem A)

German Center for Infection Research TTU 07.866 (MZ)

German Research Foundation DFG ZE 1354/2-1 (MZ)

German Research Foundation CRC1279 and CRC1310 (FK)

German Research Foundation CRC1310 (Artem A)

German Research Foundation FOR2722 (MK)

German Research Foundation FOR2830 (HH)

European Research Council ERC-StG639961 (FK)

Welch Foundation F-0003-19620604 (JSM)

Else Kröner-Fresenius-Stiftung 2024_EKEA.92 (MZ)

Stiftungsgelder 1988, KölnFortune 73/2024, CMMC CAP 41 and B11 (MZ)

## Author contributions

Conceptualization: Artem A, JAG, JSM, MZ, FK

Methodology: Artem A, JAG, VL, KH, HH, JSM, MZ, FK

Investigation: Artem A, JAG, Anna A, ZY, VL, MK, NC, LS, ZS, DS

Visualization: Artem A, JAG, VL, MZ

Formal analysis: Artem A, JAG, JSM, MZ

Funding acquisition: Artem A, MK, JSM, MZ, FK

Project administration: Artem A, JSM, MZ, FK

Supervision: JSM, MZ, FK

Writing – original draft: Artem A, JAG, MZ

Writing – review & editing: all authors

## Data and materials availability

Further information and requests for resources and reagents should be directed to and will be fulfilled by the corresponding author. Requests for materials will be fulfilled by the corresponding author upon request. Transfer of antibodies, protein constructs, and sample material might require a Material Transfer Agreement (MTA) for non-commercial usage.

Structural data are deposited at online databases and are publicly available from the date of publication:

- CryoEM structure of HCMV trimer+9 Fabs (Selection 1) – PDB ID: 9PZB, EMDB ID: EMD-72060;
- CryoEM structure of HCMV trimer+8 Fabs (Selection 2) – PDB ID: 9PZC, EMDB ID: EMD-72061.

All data that support the findings in this paper will be shared by the corresponding author upon request. Any additional information required to reanalyze the data reported in this paper is available from the corresponding author upon request.

## Supplementary Materials

### Supplementary Figures

**Fig. S1.** Characteristics of selected antibodies and cryo-EM workflow.

**Fig. S2.** Details of gH base-binding epitopes.

**Fig. S3.** Details of membrane-distal gH/gL epitopes.

**Fig. S4.** Neutralization activity of antibody cocktails against HCMV.

### Supplementary Table

**Table S1.** Cryo-electron microscopy data collection and refinement statistics.

