## Supplementary Materials for "Structural characterization of antibodies for synergistic HCMV neutralization"

**The file includes:**

Figs. S1 to S4
Table S1

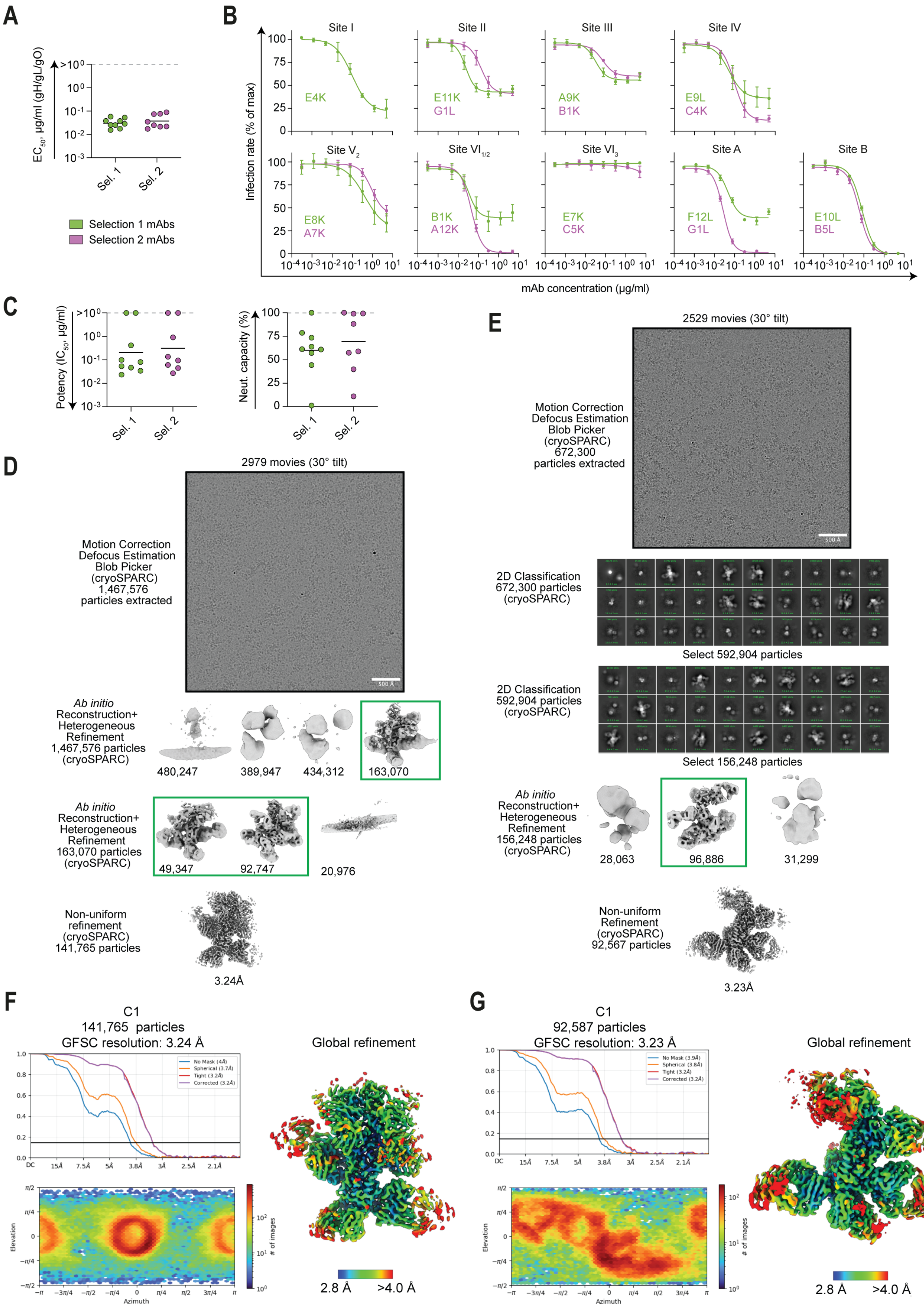


*(Caption on the next page.)*

**Fig. S1. Characteristics of selected antibodies and cryo-EM workflow.**

(**A**) Antibody binding activity with HCMV trimer measured by ELISA, represented as $EC_{50}$ values (µg/ml). Selection 1 mAbs are displayed in green, selection 2 mAbs in magenta. Each value was determined by two separate experiments, and mean values are presented. Black lines indicate the geometric mean for each selection. (**B**) Neutralization dose-response curves of mAbs acting against $HCMV_{TB40/E}$ infection in HFF-1 fibroblasts, grouped by the gH/gL binding site. Infection rates are expressed as percentages relative to the no-antibody (maximal infection) control. The mean values and SDs from three separate experiments are shown. (**C**) Neutralizing properties of mAbs against $HCMV_{TB40/E}$ infection in HFF-1 fibroblasts (from dose-response curves in **fig. S1B**). Left: Neutralization potency ($IC_{50}$, µg/ml). Right: Neutralization capacity (100% indicating complete neutralization at maximal tested antibody concentration). Each dot represents the mean of three separate experiments. Black lines indicate the geometric mean (left graph) and arithmetic mean (right graph) for each selection. (**D**) Cryo-electron microscopy data processing workflow for HCMV trimer+Selection 1 Fabs. Green boxes denote particles selected for the next round of processing. (**E**) Cryo-electron microscopy data processing workflow for HCMV trimer+Selection 2 Fabs. Green box denotes particles selected for the next round of processing. (**F**) Cryo-electron microscopy map validation for HCMV trimer+Selection 1 Fabs. Left, top shows Fourier-shell correlation (FSC) curves for the refinement. Left, bottom shows the orientation distribution plot for the refinement. The unsharpened map is shown on the right colored by estimated local resolution. (**G**) Cryo-electron microscopy map validation for HCMV trimer+Selection 2 Fabs. Left, top shows Fourier-shell correlation (FSC) curves for the refinement. Left, bottom shows the orientation distribution plot for the refinement. The unsharpened map is shown on the right colored by estimated local resolution.

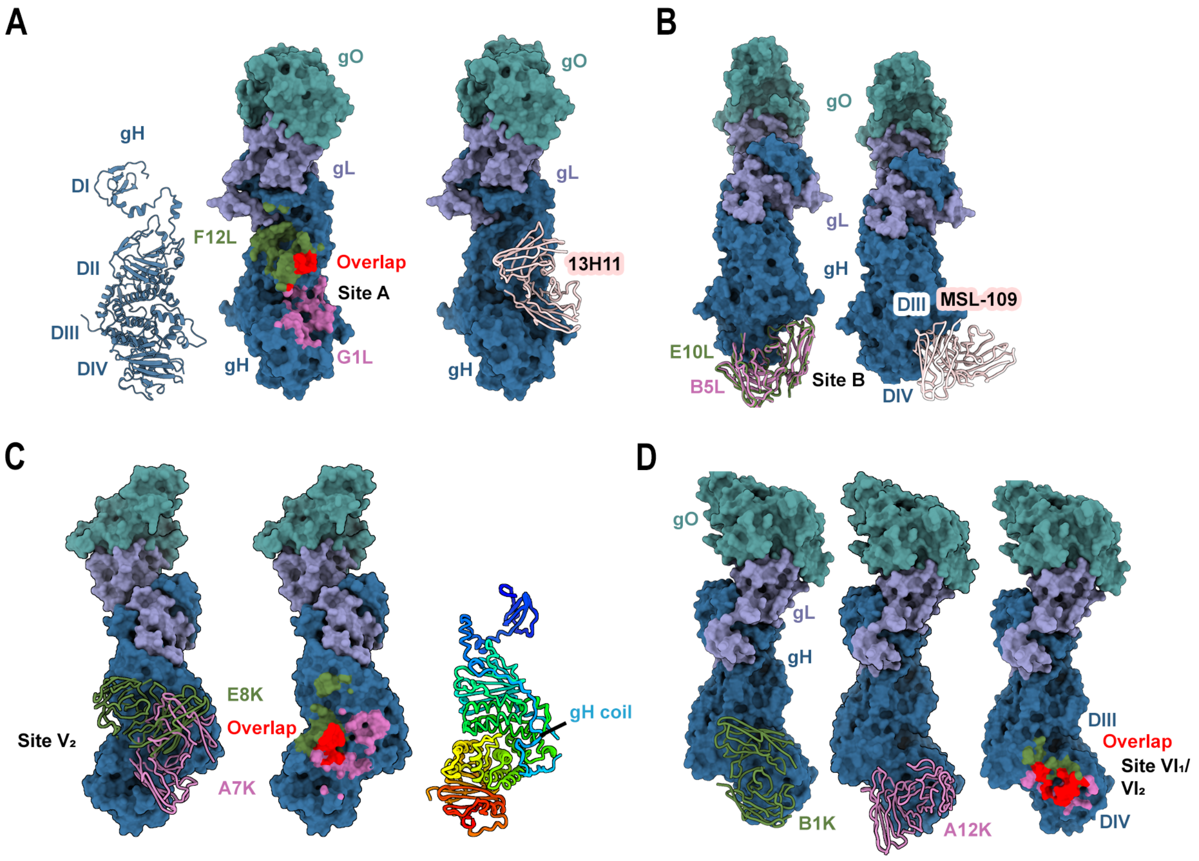


**Fig. S2. Details of gH base-binding epitopes.**

(**A**) Left: gH is shown as a ribbon with its domains DI, DII, DIII and DIV labeled. Middle: HCMV trimer is shown as a molecular surface with gH atoms within 5.5Å of F12L colored green, gH atoms within 5.5Å of G1L colored pink, gH atoms within 5.5Å both Fabs colored red. Right: HCMV trimer is shown as a molecular surface with the published antibody 13H11 shown bound to Site A as a backbone trace. (**B**) Left: HCMV trimer shown as a molecular surface with E10L (green, Selection 1) and B5L (pink, Selection 2) superimposed as backbone traces. Right: HCMV trimer is shown as a molecular surface with the published antibody MSL-109 shown as a backbone trace bound to Site B. (**C**) Left: HCMV trimer shown as a molecular surface with E8K (green, Selection 1) and A7K (pink, Selection 2) shown as backbone traces bound to Site $V_2$. Middle: HCMV trimer is shown as a molecular surface with gH atoms within 5.5Å of E8K colored green, gH atoms within 5.5Å of A7K colored pink, gH atoms within 5.5Å both Fabs colored red. Right: gH shown as a backbone trace with rainbow coloring from N- to C-terminus, and with the gH coil region labeled. (**D**) Left: HCMV trimer shown as a molecular surface with B1K shown as a backbone trace (green, Selection 1). Middle: HCMV trimer shown as a molecular surface with A12K shown as a backbone trace (pink, Selection 2). Right: HCMV trimer is shown as a molecular surface with gH atoms within 5.5Å of B1K colored green, gH atoms within 5.5Å of A12K colored pink, gH atoms within 5.5Å both Fabs colored red.

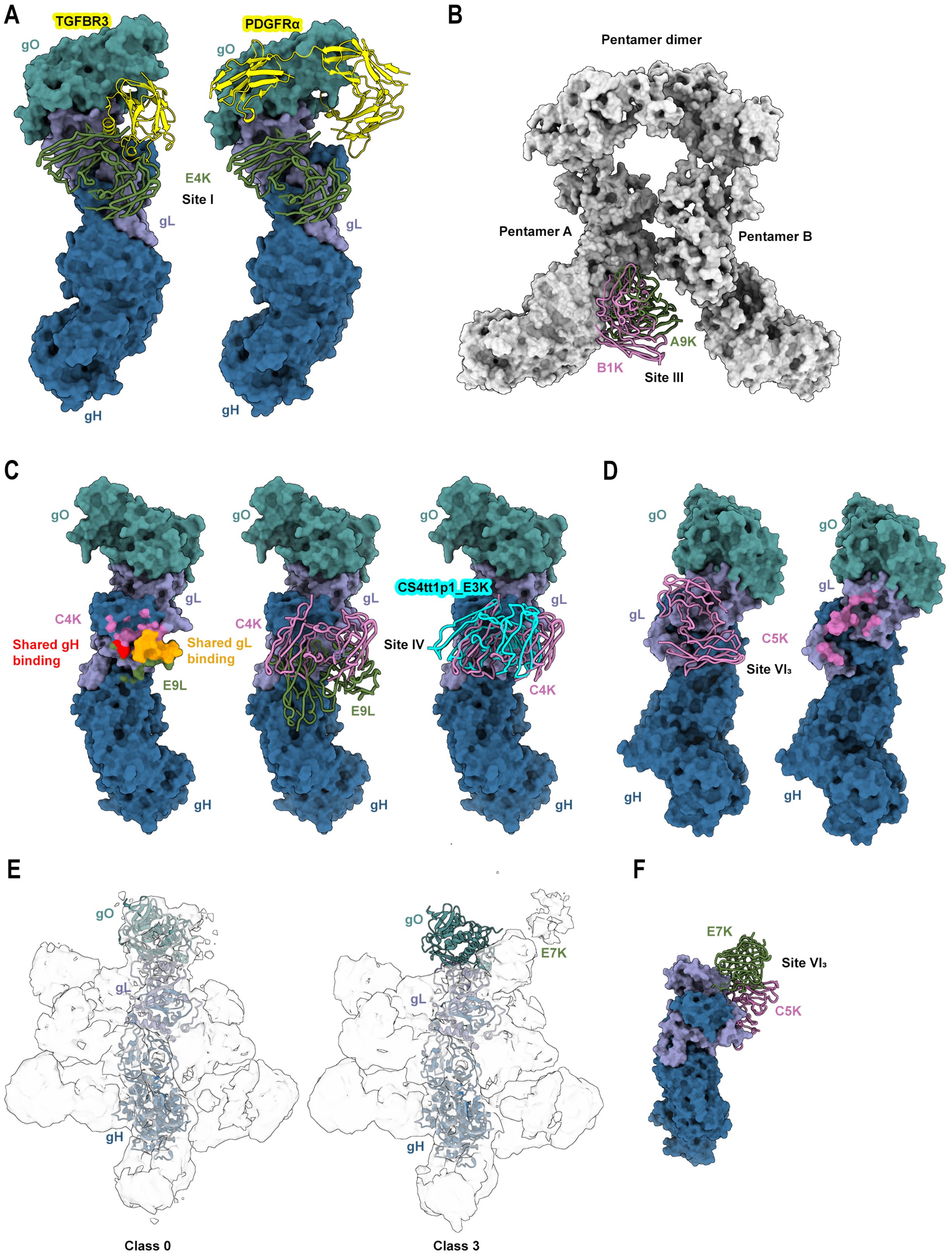


(Caption on the next page.)

**Fig. S3. Details of membrane-distal gH/gL epitopes.**

(**A**) HCMV trimer shown as a molecular surface with E4K bound to Site I (green), represented as a backbone trace and trimer receptors TGFBR3 (left) and PDGFRα (right), shown as yellow ribbons. (**B**) HCMV pentamer shown as a THBD-bound dimer (represented as a white molecular surface), with A9K and B1K backbone traces shown bound to Site III of Pentamer A in green and pink, respectively. (**C**) Left: HCMV trimer is shown as a molecular surface with gH/gL atoms within 5.5Å of E9L colored green, gH/gL atoms within 5.5Å of C4K colored pink, gH atoms within 5.5Å both Fabs colored red, and gL atoms within 5.5Å both Fabs colored orange. Middle: HCMV trimer shown as a molecular surface with E9L (green) and C4K (pink) shown as backbone traces bound to Site IV. Right: HCMV trimer shown as a molecular surface with C4K (pink) and the previously published Fab CS4tt1p1_E3K (cyan) shown as backbone traces bound to Site IV. (**D**) HCMV trimer showed as a molecular surface with C5K (pink) bound to site $VI_3$. On the left, C5K is shown as a backbone trace and on the right, HCMV trimer atoms within 5.5Å of C5K are colored pink. (**E**) Left: reconstructed cryo-EM map of a 3D class containing gO density but no E7K. Right: reconstructed cryo-EM map of a 3D class containing E7K bound in the usual position of gO. HCMV trimer containing gO is shown as backbone trace in both maps. (**F**) Superposition of E7K modeled into Class 3 from **fig. S3E** and C5K at site $VI_3$ showing their steric clash.

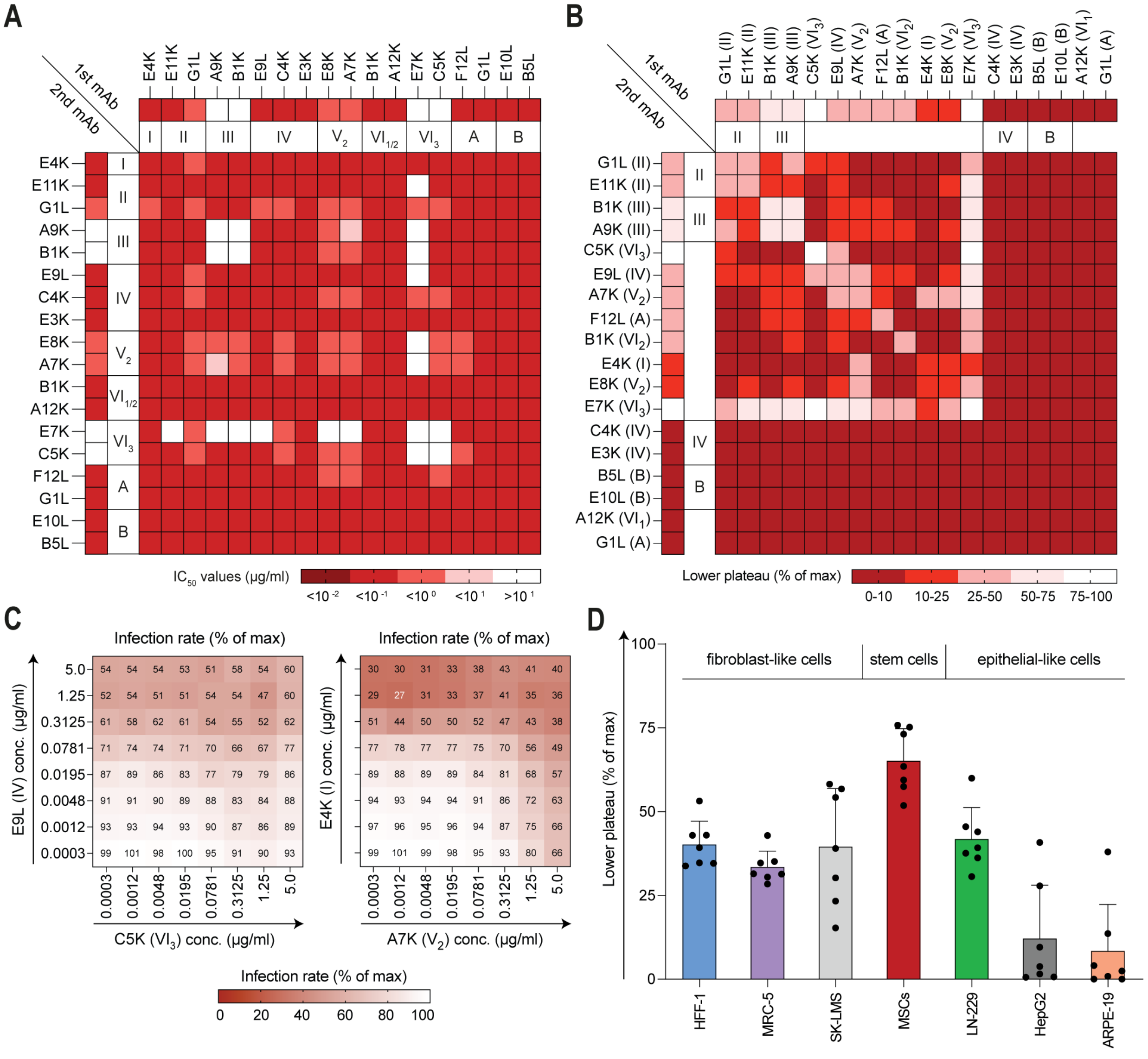


*(Caption on the next page.)*

**Fig. S4. Neutralization activity of antibody cocktails against HCMV.**

(**A**) Neutralizing performance of individual mAbs (top and left panels) and their cocktails (heatmap) in blocking $HCMV_{TB40/E}$ infection in HFF-1 fibroblasts, represented as $IC_{50}$ values (µg/ml) in sequential red colors. The graph is mirrored along the main diagonal. mAbs are clustered by epitope representation. (**B**) Lower plateau values (i.e. residual infection levels) of individual mAbs (top and left panels) and their cocktails (heatmap) against $HCMV_{TB40/E}$ infection in HFF-1 fibroblasts, displayed in sequential red colors. The graph is mirrored along the main diagonal. mAbs are clustered by additive neutralization levels and epitope representation (see also **Fig. 4B**). 'High-plateau' mAbs and their combinations were tested in at least two separate experiments; representative data are shown. (**C**) Neutralizing activity of non-synergistic cocktails with different stoichiometries assessed by testing varying concentrations of cocktail members against $HCMV_{TB40/E}$ infection in HFF-1 fibroblasts. Percentage of residual infection achieved by each concentration is shown. Mean values from two separate experiments are displayed. (**D**) Average lower plateau levels of the tested 'high-plateau' mAbs for different host cell lines. The mAbs included in the analysis are: E4K (I), E11K (II), A9K (III), E9L (IV), A7K ($V_2$), B1K ($VI_2$), F12L (A). Primary data are sourced from **Fig. 4F**, HFF-1 data of $HCMV_{TB40/E}$ infection from **Fig. 4E**. Error bars represent SDs. Each dot represents the result for an individual mAb, showing the mean from two separate experiments.

| EM data collection | HCMV trimer+9 Fabs (Selection 1) | HCMV trimer+8 Fabs (Selection 2) |
|---|---|---|
| Microscope | Glacios | Glacios |
| Detector | Falcon 4 | Falcon 4 |
| Defocus range (μm) | 0.8–2.2 | 0.8–2.2 |
| Pixel size (Å) | 0.94 | 0.94 |
| Tilt angle (°) | 30 | 30 |
| Magnification | 150,000x | 150,000x |
| Voltage (kV) | 200 | 200 |
| Exposure rate ($e^-$/pix/sec) | 2.9 | 2.9 |
| Micrographs collected | 2,979 | 2,529 |
| Micrographs used | 2,445 | 1,306 |
| Electron exposure ($e^-$/Å$^2$) | 50 | 50 |
| Particles extracted | 1,467,576 | 672,300 |
| Automation software | SerialEM | SerialEM |
| **3D reconstruction statistics** | | |
| Final particles | 141,765 | 92,567 |
| Symmetry | C1 | C1 |
| Unmasked resolution at 0.5 FSC (Å) | 7.7 | 7.7 |
| Masked resolution at 0.5 FSC (Å) | 3.7 | 3.7 |
| Unmasked resolution at 0.143 FSC (Å) | 4.0 | 3.9 |
| Masked resolution at 0.143 FSC (Å) | 3.2 | 3.2 |
| **Model refinement and validation** | | |
| Refinement package | Phenix | Phenix |
| Refinement tool | Real-space refinement | Real-space refinement |
| Refinement strategies | min global, local_grid_search, adp, ss restraints, rotamer restraints, Ramachandran restraints | min global, local_grid_search, adp, ss restraints, rotamer restraints, Ramachandran restraints |
| Initial model(s) used (PDB ID) | 7LBE | 7LBE |
| Model composition | | |
| Number of atoms | 19,420 | 19,426 |
| Protein B-factors (mean) | 82.5 | 99.6 |
| RMSD bond lengths (Å) | 0.646 | 0.619 |
| RMSD bond angles (°) | 0.003 | 0.004 |
| MolProbity score | 1.33 | 1.48 |
| Clashscore | 3.76 | 5.1 |

| | | |
|---|---|---|
| Rotamer outliers (%) | 0.0 | 0.1 |
| C-beta outliers (%) | 0.0 | 0.0 |
| Ramachandran plot | | |
| Favored (%) | 97.0 | 96.7 |
| Allowed (%) | 3.0 | 3.3 |
| Outliers (%) | 0 | 0 |
| EMRinger score | 4.49 | 3.89 |
| CaBLAM outliers (%) | 1.9 | 1.7 |
| CC (mask) | 0.83 | 0.85 |
| **Data Availability** | | |
| EMDB | EMD-72060 | EMD-72061 |
| PDB | 9PZB | 9PZC |

**Table S1. Cryo-electron microscopy data collection and refinement statistics.** Statistics for data collection, processing, model building, and refinement are shown.
